# Jump-start and push-start: Mutual activation through crosstalk in coupled quorum sensing pathways

**DOI:** 10.1101/2023.05.08.539885

**Authors:** Joseph Sanders, Hoda Akl, Stephen J. Hagen, BingKan Xue

## Abstract

Many bacterial species are able to coordinate population-wide phenotypic responses through the exchange of diffusible chemical signals, a behavior known as quorum sensing. A quorum sensing bacterium may employ multiple types of chemical signals and detect them using inter-connected pathways that crosstalk with each other. While there are many hypotheses for the advantages of sensing multiple signals, the prevalence and functional significance of crosstalk between the sensing pathways are much less understood. Here we explore the effect of intra-cellular signal crosstalk on a simple model of a quorum sensing circuit. The model captures key aspects of typical quorum sensing pathways, including detection of multiple signals that crosstalk at the receptor and promoter levels, positive feedback, and hierarchical positioning of sensing pathways. We find that a variety of behaviors can be tuned by modifying crosstalk and feedback strengths. These include activation or inhibition of one output by the non-cognate signal, broadening of dynamic range of the outputs, and the ability of either the upstream or downstream branch to modulate the feedback circuit of the other branch. Our findings show how crosstalk between quorum sensing pathways can be viewed not solely as a detriment to the flow of information but also as a mechanism that enhances the functional range of the full regulatory system: When positive feedback systems are coupled through crosstalk, several new modes of activation or deactivation become possible.

## 1 Introduction

Many bacteria regulate and synchronize population-wide behaviors by exchanging diffusible chemical signals with other individuals of the same or different species within the community [1]. By secreting these chemical signals, known as autoinducers, and detecting their local concentrations, the bacteria can induce phenotypes collectively, in response to environmental and population conditions. These quorum-sensing regulatory pathways are usually sensitive to a variety of environmental and cross-species cues in addition to their own autoinducers, so that they control multiple phenotypic outputs in a complex fashion [2].

Quorum sensing bacteria typically synthesize and detect more than one chemically distinct autoinducer, often with positive feedback controlling the rate of autoinducer production. The different autoinducers are detected by cognate receptors that drive regulatory pathways coupled to varying degrees [3]. The ability to sense more than one autoinducer is hypothesized to provide a number of potential benefits to a microbial species. It may offer advantages in interspecies interactions, including greater resistance to manipulation by other species [4] or the ability for both interspecies and intraspecies communication [5]. Multiple signals may also allow temporal control of distinct phenotypes if different autoinducers accumulate at different rates [6, 7], or they may help infer physical conditions such as spatial confinement [8], or provide advantages in quorum cheating [9]. Sensing through multiple signals and receptors in general may allow more sophisticated control of output dynamics of a sensing pathway [10]. But the ligand-specificity of quorum sensing receptors varies considerably among species and strains [4]; autoinducers employed by one organism often elicit a response from non-cognate receptor pathways in related variants or other microbial species. The lack of signal specificity allows interspecies “crosstalk” in bacterial communities [11], a phenomenon that has been widely explored in the context of social behaviors such as kin discrimination, eavesdropping and facultative cheating [3].

Crosstalk can also occur within a single species or strain. A pathway that senses one autoinducer may also be activated or inhibited by other autoinducers produced by the same organism. As these effects may occur through several different mechanisms, several definitions of crosstalk have arisen. In a system of two signal/receptor pairs that drive different promoters, and where specificity is poor, crosstalk has been characterized in terms of whether the lack of specificity resides in the ligand/receptor interactions or at the promoter-binding level [12]. It is also common, however, for pathways that detect multiple signals to funnel down to a fewer number of downstream outputs [10]. An extreme case is *Vibrio harveyi*, which senses three distinct autoinducers, each with its own dedicated sensor kinase; information from the three kinases is funneled into control of the same phosphorelay system [13]. Such funneled architecture has been described as crosstalk [14]. Here, we find it useful to define crosstalk as any mixing between two signaling pathways A and B that have their own signal inputs and outputs, but wherein signal A also modulates to some extent the output of receptor B, and vice versa. Crosstalk is a degree of coupling between two functional sensing pathways in the same organism [15, 16, 17], and the strength of crosstalk lies along on a continuum from very strong (funneled) to very weak (orthogonal pathways).

Because crosstalk mixes information received from separate signals, it would appear likely to degrade the performance of a quorum sensing pathway. It is however a highly evolvable property that can be reduced or even eliminated through (for example) receptor design [18, 19, 4]. Therefore, although there exist several hypotheses for why bacterial species use multiple autoinducer signals, there is still little understanding of why crosstalk is common in quorum sensing systems, and how it affects the output behaviors of these networks, beginning at the level of an individual organism. Gram negative quorum sensing systems that employ autoinducers of the acyl homoserine lactone (HSL) type are particularly susceptible to crosstalk, as the HSLs are chemically similar and their cognate receptors typically respond to HSLs spanning a range of acyl chain lengths [11]. The quorum sensing system in the bacterium *Vibrio fischeri* is a model example [20] with homologs in numerous other species [4]. We will focus on two pathways in that organism, LuxI/R and AinS/R, which are subject to several forms of crosstalk, shown in Figure 1. The *lux* operon that controls bioluminescence is under immediate control of the LuxI/R pathway, a feedback loop in which LuxI is the synthase for the autoinducer 3-oxo-C6-homoserine lactone (3OC6-HSL) that interacts with the intracellular receptor LuxR to bind the *lux* promoter. However, the production of LuxR is modulated by LitR, which is controlled by a second, upstream quorum sensing pathway, AinS/R. AinS and AinR synthesize and detect respectively an autoinducer N-octanoyl-L-homoserine lactone (C8-HSL) to control LitR production. The LuxI/R and AinS/R pathways crosstalk through several mechanisms. The LuxR-3OC6-HSL complex is able to modulate expression of AinS, which encodes the C8-HSL synthase, by interacting with a *lux* -box-type binding site [21, 20]. In addition, C8-HSL can also interact with LuxR to promote its binding to the *lux* -box. Thus, C8-HSL exerts an influence on the downstream LuxR/LuxI system via LuxR and LitR, while 3OC6-HSL influences the production of the C8-HSL synthase upstream.

**Figure 1:**
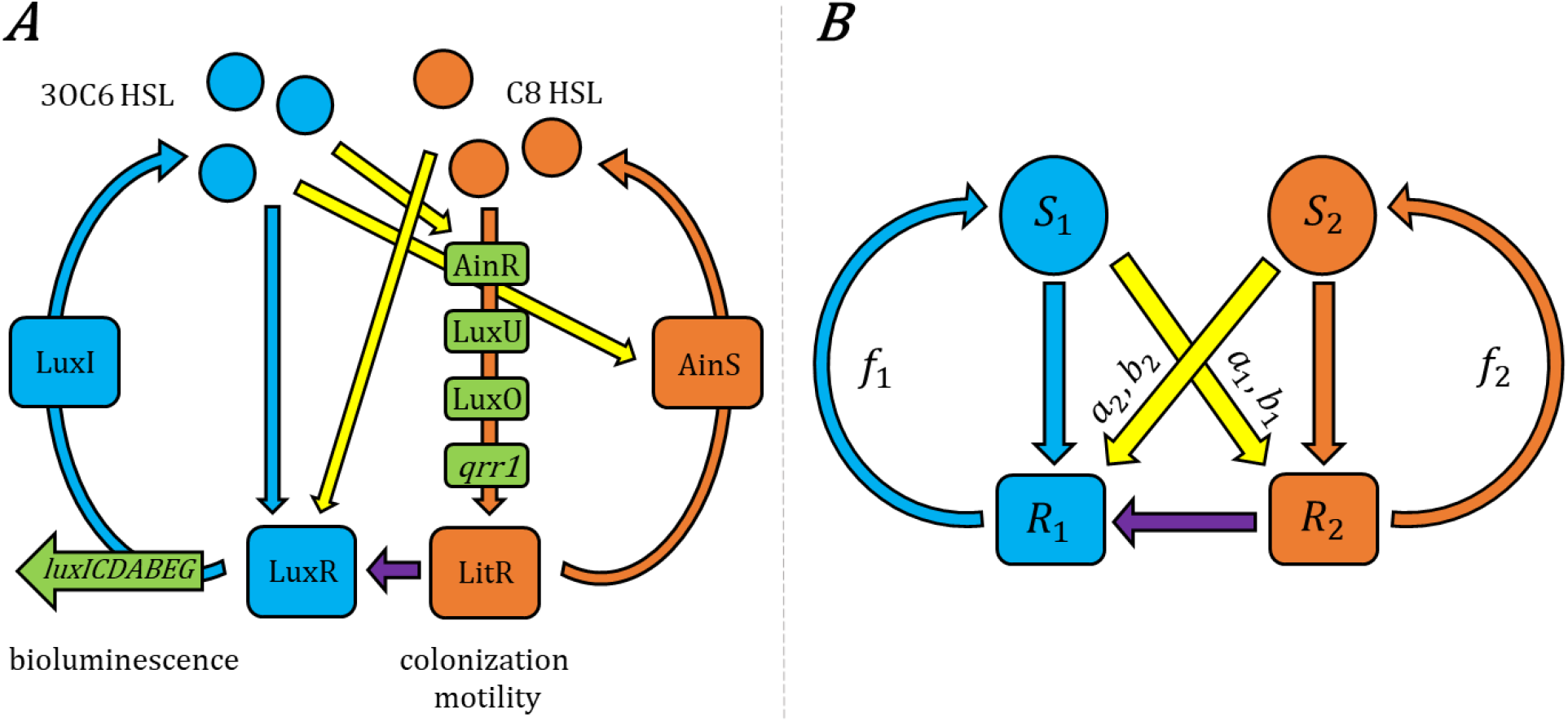
(A) The AinR/AinS and LuxR/LuxI quorum sensing pathways in *Vibrio fischeri*, which primarily control colonization traits and bioluminescence respectively, interact through several crosstalk mechanisms [22, 20]. (A third pathway involving LuxS, LuxP/LuxQ and the autoinducer AI-2, coupled to the above through LuxU/LuxO, is not shown here.) The histidine kinase AinR detects its cognate autoinducer C8-HSL produced by AinS, initiating the LuxU/LuxO phosphorylation pathway. This pathway controls the expression of the regulatory RNA *qrr1*, a post-translational repressor of *litR*. In addition to controlling phenotypes related to motility and host colonization, LitR modulates production of LuxR, which is the intracellular receptor for the autoinducer 3OC6-HSL of the LuxI/LuxR pathway. LuxR becomes a transcriptional activator for the *lux* operon when bound either to its cognate signal 3OC6-HSL, produced by LuxI, or to the non-cognate C8-HSL. In addition, 3OC6-HSL interacts with LuxR to modulate activation of *ainRS*. Thus the AinR/AinS and LuxR/LuxI pathways both respond to each others’ autoinducers, while the AinR/AinS pathway also acts upstream of the LuxR/LuxI pathway through LitR. (B) The simplified model studied in this work captures the key elements of a quorum sensing system with crosstalk: Two signals (*S*_1_, *S*_2_) elicit their respective cognate responses (*R*_1_, *R*_2_), with crosstalk between them (yellow arrows) and an additional link between *R*_1_ and *R*_2_ that makes *R*_2_ upstream of *R*_1_. The signals are produced with positive feedback (*f*_1_, *f*_2_) from their respective responses. The crosstalk parameters *b*_*i*_ and *a*_*i*_ respectively define the strength of cross-binding (between each signal *S*_*i*_ and its non-cognate receptor) and cross-activation (of the non-cognate response); see Appendix A.1.

In order to understand how the tuning of crosstalk strength affects a two-pathway quorum sensing system, we have analyzed a simplified model of the *V. fischeri* system. The model retains key features of two signals that primarily stimulate two responses, with crosstalk as well as positive feedback in autoinducer synthesis. We use this streamlined model to explore how different aspects of the crosstalk interact with feedback to reshape the steady state outputs that are available to the system.

## 2 Model

We consider a model represented by the schematic in Figure 1B, capturing the essential elements of crosstalk in the AinR/AinS and LuxR/LuxI quorum sensing pathways of *V. fischeri*. There are two signaling pathways, each of which produces a signal (*S*_1_ or *S*_2_) that induces a response (*R*_1_ or *R*_2_, respectively). The signal associated with each pathway is produced with positive feedback (*f*_1_, *f*_2_) from the response. One pathway (*R*_2_, *S*_2_) is functionally “upstream” of the other (*R*_1_, *S*_1_), in that response *R*_1_ is dependent on *R*_2_ (we do not call this link “crosstalk” because *R*_1_ cannot function without it, so this link is not a tunable perturbation, see discussion in Section 4.1). In addition, each of the two signals has some effect on the response of the other (“non-cognate”) pathway.

In the steady state, the elements of multiple signals, feedback and crosstalk are captured by the equations:

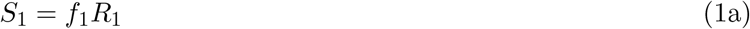

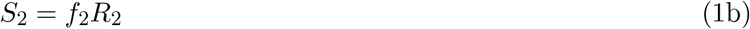

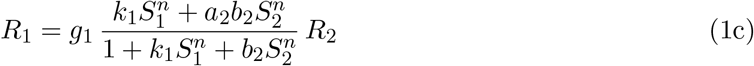

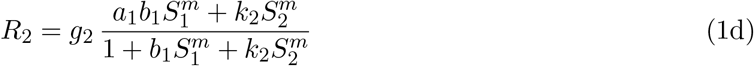

The origin of these equations in the steady state output of a two-pathway sensing system is explained in Appendix A.1. In these equations, *S*_1_ and *S*_2_ represent the concentrations of the two autoinducers; *R*_1_ and *R*_2_ represent the expression levels of the quorum-regulated genes of each pathway, including genes that encode the autoinducer synthases. Eqs. (1a, 1b) relate the signal concentrations to the regulated genes, due to positive feedback. Eqs. (1c, 1d) give the steady state response to the two signal levels. Among the parameters, *f*_*i*_ is the feedback strength for each autoinducer, which depends on the rate of signal synthesis and the population density of cells; *g*_*i*_ is the maximum expression level of *R*_*i*_; *k*_*i*_ is the interaction strength of a receptor with its cognate signal. For noncognate (crosstalk) interactions, the strength of binding and activation are described separately: *b*_*i*_ (“binding”) captures the ability of a signal to competitively bind its non-cognate receptor; *a*_*i*_ (“activation”) describes the efficiency of a signal, when bound to the non-cognate receptor, in cross-activating the non-cognate response. Finally, the exponents *n* and *m* represent the cooperativity of the signal response in each pathway.

We can simplify the equations by rescaling *S*_1_ and *S*_2_ to set *k*_1_ = *k*_2_ = 1, and rescaling *R*_1_ and *R*_2_ to set *g*_1_ = *g*_2_ = 1 (see Appendix A.1). Then the maximum value for the rescaled *R*_1_ and *R*_2_ is 1 (for *a*_*i*_ ≤ 1). Of the remaining parameters, *a*_*i*_ and *b*_*i*_ control the crosstalk strength. If the binding *b*_*i*_ = 0, then the signal from pathway *i* cannot elicit any response from the non-cognate pathway, so there is no crosstalk. On the other hand, if the activation *a*_*i*_ = 0, the signal from pathway *i* may bind to but not activate the response of the non-cognate pathway. For *a*_*i*_ *>* 0 there is potential cross-activation between the pathways; we assume *a*_*i*_ ≤ 1, which means the cross-activation by the non-cognate signal cannot be more efficient than the cognate signal. In most of what follows, we focus on the parameters *a*_2_ and *b*_1_, and set the other two parameters *a*_1_ = *b*_2_ = 1. That is, we focus on the case where *S*_2_ interacts strongly with the noncognate receptor, but the complex is not necessarily an efficient activator for *R*_1_. This is partly motivated by the example of *V. fischeri*, in which the effect of C8-HSL (*S*_2_) on *lux* expression (analogous to *R*_1_) is well known [23], and 3OC6-HSL (analogous to *S*_1_) has been shown to stimulate the *ainRS* pathway (*R*_2_) [21]. We will further assume that *m* = *n*, and consider a range of values for the cooperativity *n*.

The responses *R*_1_ and *R*_2_ are not simply functions of *S*_1_ and *S*_2_ as in Eqs. (1c,1d), except in the special circumstance where signal levels are externally controlled. In natural settings the signals are tied to the responses through the feedback *f*_1_ and *f*_2_. As a result, the equilibrium values of all signals and responses are determined by the feedback strengths by solving Eqs. (1a–1d). Eliminating *S*_1_ and *S*_2_ from Eqs. (1c,1d) using Eqs. (1a,1b) and simplifying the parameters as described above, we arrive at two self-consistent equations for *R*_1_ and *R*_2_:

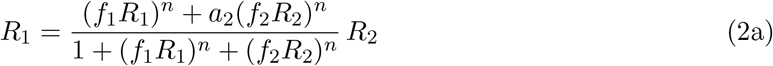

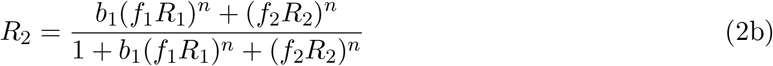

We find the solutions to these equations using numerical solvers from the SciPy package for Python. We first solve a set of ODEs for which Eqs. (2) are the equilibrium (see Appendix A.2). We integrate these ODEs for a sufficient amount of time that the variables come close to an equilibrium. Then, we use these values as initial guesses for a root solver to find the precise solutions. This method allows us to find multiple solutions if they exist and are stable, by using many random initial values in solving the ODEs. Once the solutions are refined using the root solver, we remove redundant solutions that have already been found (see Appendix A.2 for details).

## 3 Results

The solutions to our main equations (2a,2b) represent the responses *R*_1_ and *R*_2_ as functions of the feedback strengths *f*_1_ and *f*_2_. Increased feedback strength may correspond to the condition of high cell density, where the autoinducer is captured by neighboring cells rather than being lost to the environment. When the cell density reaches a certain level, a phenotypic response is triggered. Our goal is to see how this response is modulated by the crosstalk parameters *a*_2_ and *b*_1_. Without crosstalk, the pathways operate independently: response *R*_*i*_ will be activated if the feedback *f*_*i*_ reaches a certain level. Crosstalk allows the feedback *f*_1_ not only to elicit the cognate response *R*_1_ but also to influence the other response *R*_2_, and vice versa. We will characterize such effects below.

### 3.1 Crosstalk can both activate and inhibit non-cognate responses

Figure 2 shows a heat map of *R*_1_ as a function of *f*_1_ and *f*_2_. *R*_1_ is of interest as it is the most downstream element in our circuit (Figure 1B), affected by both signals and the upstream response *R*_2_. When both crosstalk parameters are at full strength, *a*_2_ = *b*_1_ = 1 (given that we also assume *a*_1_ = *b*_2_ = 1), *R*_1_ is determined simply as a linear combination of *f*_1_ and *f*_2_ (Figure 2 bottom-right panel). It means that *R*_1_ is equally well activated by either one of the quorum signals, either directly by the cognate *S*_1_ or through crosstalk by the non-cognate *S*_2_. This strong crosstalk is analogous to what occurs in *V. harveyi* quorum sensing network, where three autoinducer inputs add linearly to give a single output [24].

**Figure 2:**
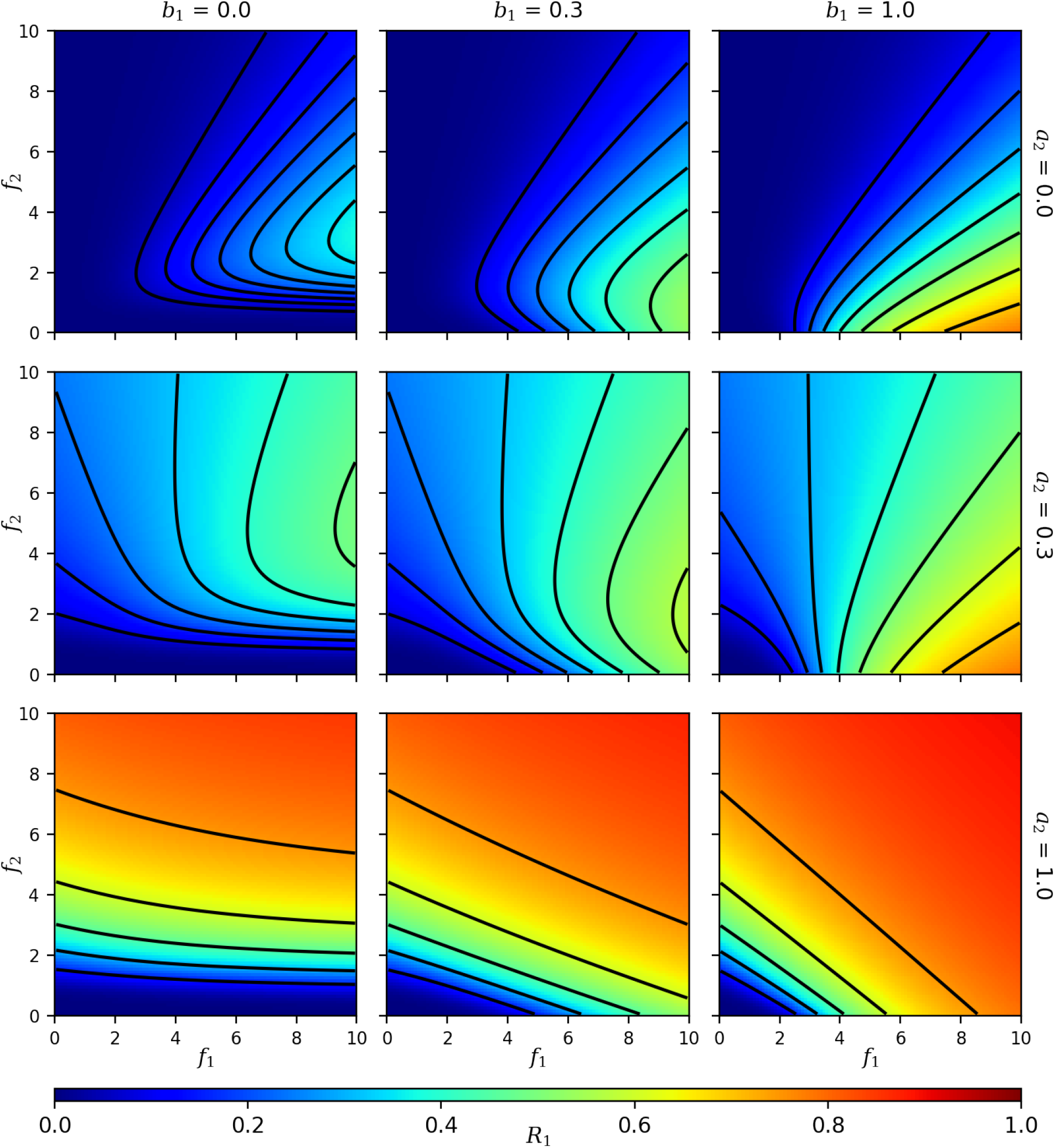
Heat maps showing the value of *R*_1_ as a function of the feedback strengths *f*_1_ and *f*_2_. Rows correspond to different values of the downstream-directed crosstalk activation strength *a*_2_, whereas columns correspond to values of the upstream-directed crosstalk binding strength *b*_1_ (all panels have *a*_1_ = *b*_2_ = 1, *n* = 1). Black lines are contours of constant *R*_1_.

Figure 2 also shows how the strength of the crosstalk *a*_2_ determines whether the non-cognate signal *S*_2_ activates or inhibits the response *R*_1_. Strong activating crosstalk *a*_2_ (Figure 2 bottom row) allows *R*_1_ to increase with *f*_2_. Weakly activating crosstalk, where *a*_2_ is small (Figure 2 top row), allows a large *f*_2_ to inhibit *R*_1_. This is because at the low *a*_2_ limit cross-activation is inefficient, so that non-cognate binding (*b*_2_ = 1) allows competitive inhibition of *R*_1_ by *S*_2_.

Another feature to note is that, when the crosstalk binding strength *b*_1_ is very small (Figure 2 left column) and *f*_2_ is also small, no amount of *f*_1_ can activate *R*_1_. This is because *R*_1_ relies on the upstream response *R*_2_, which remains off under conditions of small *f*_2_ and *b*_1_. However, as *b*_1_ increases (right column), part of the small-*f*_2_ region can now be activated by *f*_1_ alone: Crosstalk from the downstream signal *S*_1_ to the upstream response *R*_2_ can activate the downstream response *R*_1_. In Section 3.3 below we elaborate on this mechanism where crosstalk from the downstream signal activates both responses.

### 3.2 Crosstalk can modulate the dynamic range of joint responses

To study the dynamic range of both responses *R*_1_ and *R*_2_ together, we make a parametric plot of their values as the feedback *f*_1_ and *f*_2_ are varied (Figure 3). This creates a mesh of possible solutions that deforms as the crosstalk strengths *a*_2_ and *b*_1_ change. The mesh lies in the upper left half of each panel, because *R*_1_ ≤ *R*_2_ as a result of Eq. (2a): the downstream response *R*_1_ relies on the upstream *R*_2_. *R*_1_ and *R*_2_ show greatest range and span a broader, two-dimensional region of the graph when crosstalk is weak, i.e., with *a*_2_ small (Figure 3 top row). As the crosstalk strength increases, the *R*_1_, *R*_2_ responses become more tightly coupled and span a reduced area. The effect is most apparent when both *a*_2_ and *b*_1_ approach 1 (Figure 3 bottom right), for which the 2D mesh collapses toward a single curve. *R*_1_ and *R*_2_ are then tied together, and are both linear in *f*_1_ and *f*_2_ as seen from Figure 2 (bottom right; see also Figure S1).

**Figure 3:**
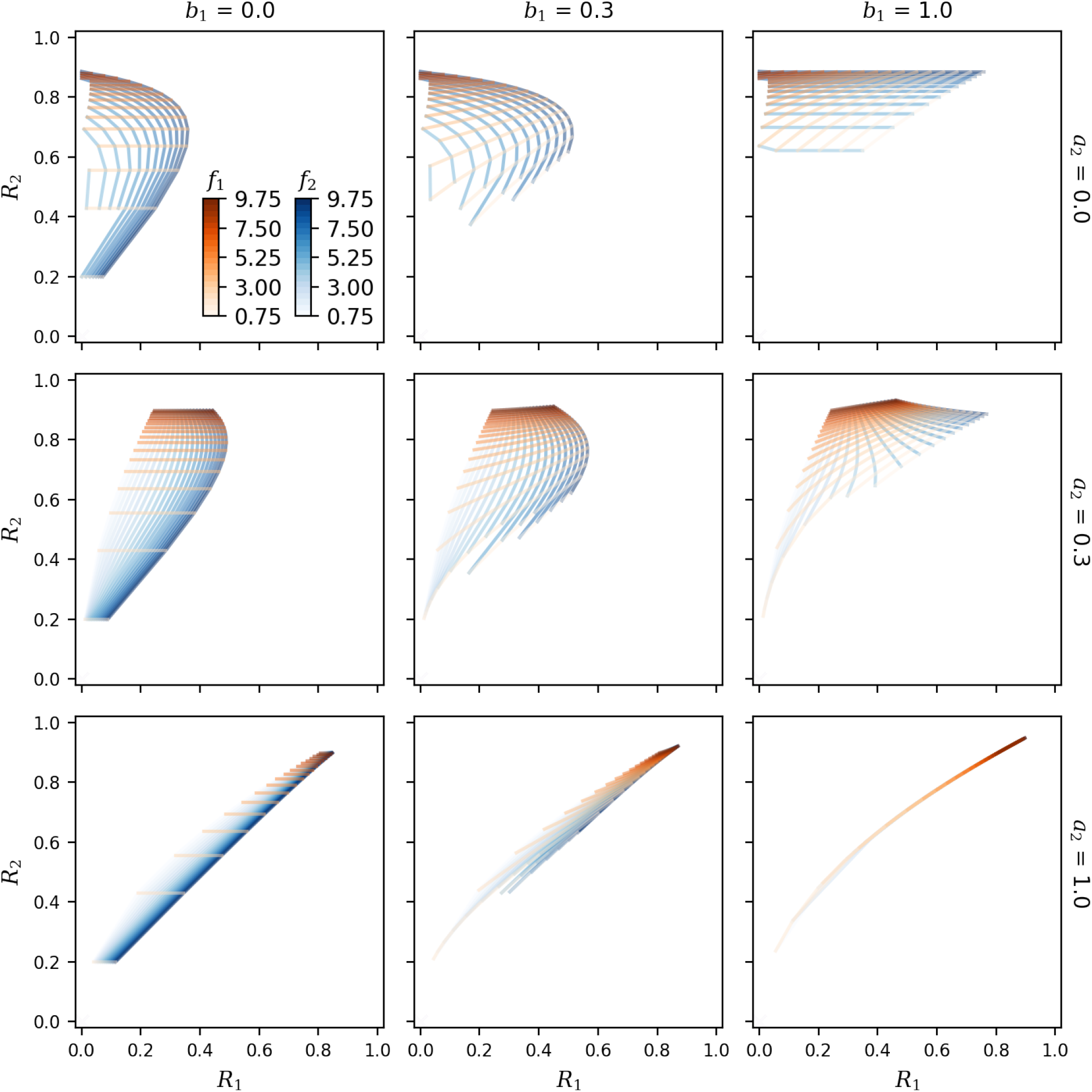
Mesh plots showing the steady state *R*_1_ and *R*_2_ at different feedback strengths. Orange curves represent constant *f*_1_ values, and blue curves represent constant *f*_2_ values. Rows and columns correspond to different values of the crosstalk parameters *a*_2_ and *b*_1_, respectively (with *a*_1_ = *b*_2_ = 1, *n* = 1).

### 3.3 Crosstalk can facilitate new mechanisms of activating responses

To visualize how both responses *R*_1_ and *R*_2_ depend on the feedback *f*_1_ and *f*_2_, we create “ellipse plots” in Figure 4, in which an array of ellipses displays the values of both *R*_1_ and *R*_2_ at different positions in the (*f*_1_, *f*_2_) plane. For each ellipse, the horizontal axis is proportional to the value of *R*_1_, and the vertical axis is proportional to *R*_2_. Thus, the width of each ellipse in each panel of Figure 4, as a function of *f*_1_ and *f*_2_, matches the *R*_1_ value shown in the heat map of Figure 2, while the height of each ellipse represents *R*_2_ (Figure S1). Where there are multiple stable solutions at the same (*f*_1_, *f*_2_) point, we overlay multiple ellipses on top of one another. In particular, a small black dot (a vanishing ellipse) indicates that the trivial solution *R*_1_ = *R*_2_ = 0 is stable.

**Figure 4:**
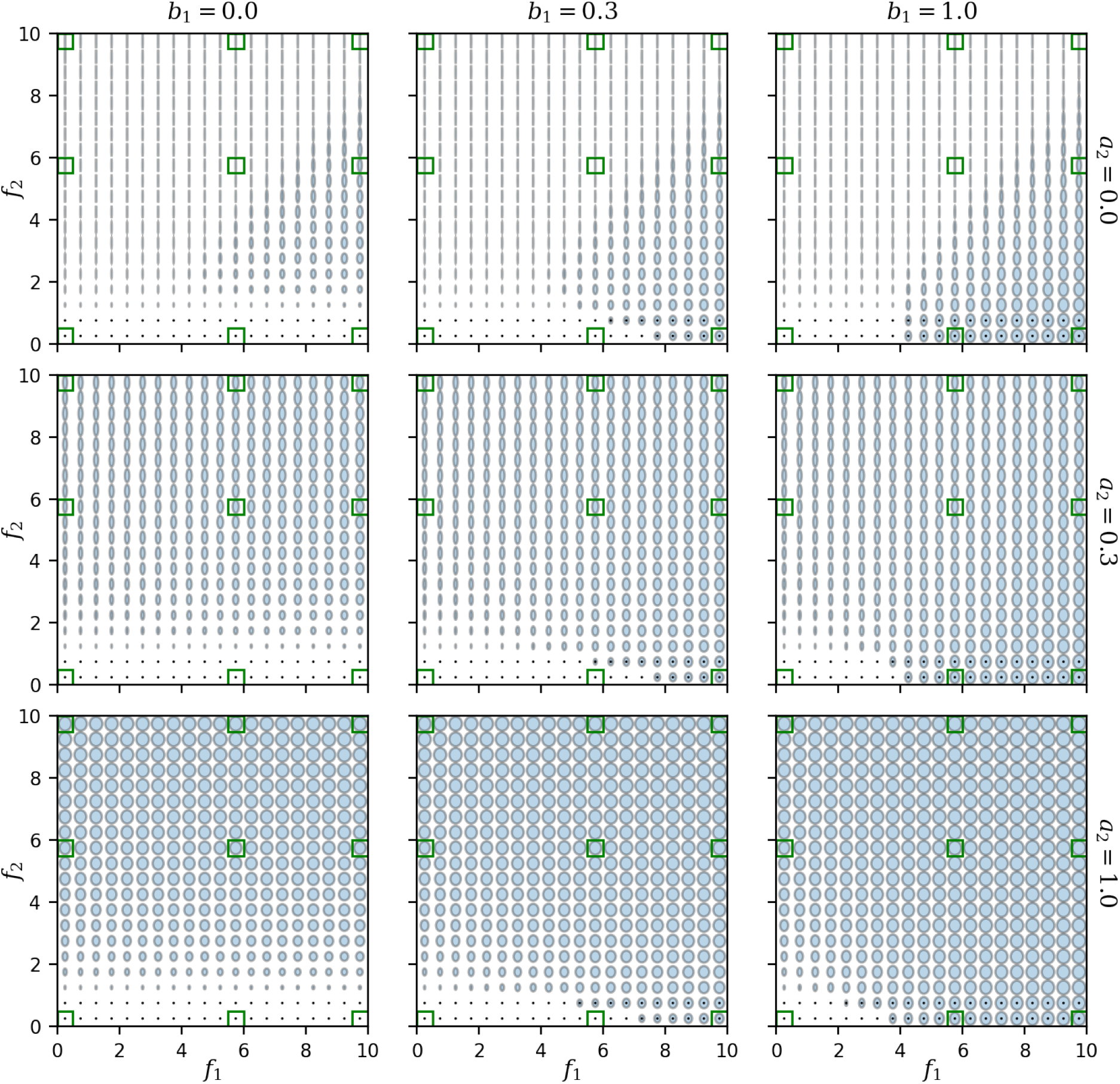
“Ellipse plots” showing the responses *R*_1_ and *R*_2_ simultaneously as functions of the feedback strengths *f*_1_ and *f*_2_. The width and height of each ellipse represent the values of *R*_1_ and *R*_2_, respectively, at a given point in the (*f*_1_, *f*_2_) plane. A black point superimposed on an ellipse indicates that the trivial state *R*_1_ = *R*_2_ = 0 is also stable. Rows and columns represent different values of the crosstalk parameters *a*_2_ and *b*_1_, respectively (with *a*_1_ = *b*_2_ = 1, *n* = 1). The *f*_1_ and *f*_2_ values marked in green are further explored in Figure 5.

The ellipse plots of Figures 4 contain all the information in our results. As with the heat map in Figure 2 and the mesh plots in Figure 3, we see that when *a*_2_ increases (from top to bottom rows), the upper left part of each graph shows a stronger *R*_1_ response. In addition, from the ellipse plots in Figure 4 it is clear that when the upstream crosstalk *b*_1_ increases, the lower right part of each graph with a small feedback *f*_2_ changes from having no response to having both *R*_1_ and *R*_2_ activated.

Figure 5 presents a different slice through the parameter space by showing *R*_1_ and *R*_2_ as functions of *a*_2_ and *b*_1_ in each panel, for selected values of *f*_1_ and *f*_2_ which vary between the panels. (The *f*_1_ and *f*_2_ values used for these plots are indicated by green squares in Figure 4.) Thus, each panel allows us to move across the different panels in Figure 4, showing how the crosstalk strengths change the responses at fixed feedback strengths. For a large *f*_2_ and relatively small *f*_1_ (Figure 5 top left), the *R*_1_ response can be activated by increasing *a*_2_ even though *R*_1_’s cognate feedback *f*_1_ is weak. Similarly, in the case of high *f*_1_ and low *f*_2_ (Figure 5 bottom right), increasing *b*_1_ will activate not only the response *R*_1_ as expected from a high *f*_1_, but also the response *R*_2_ despite weak *f*_2_.

**Figure 5:**
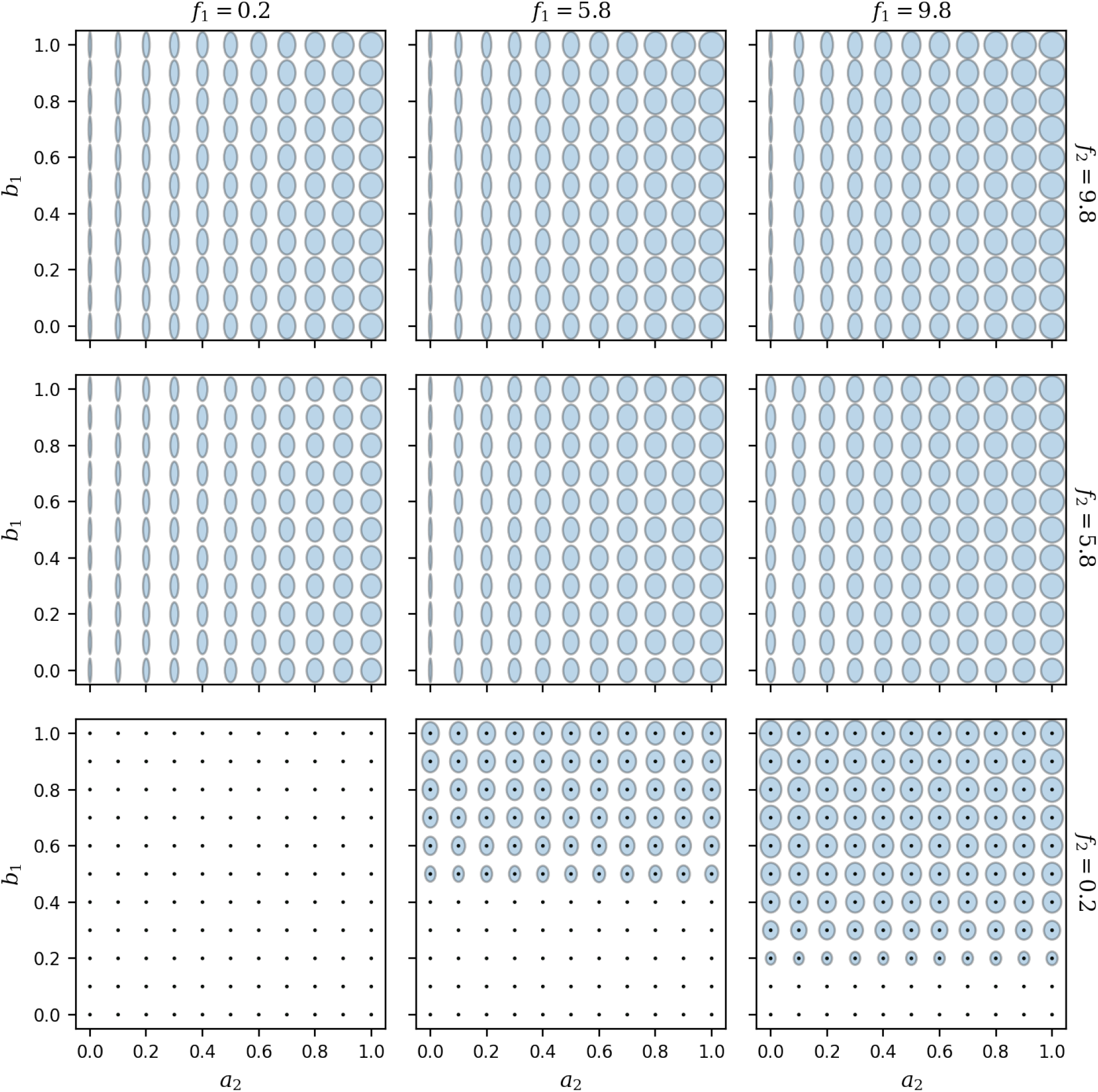
Dependence of the responses *R*_1_ and *R*_2_ on the crosstalk parameters *a*_2_ and *b*_1_. With the same parameters as in Figure 4, the width and height of each ellipse represent the values of *R*_1_ and *R*_2_, respectively. Rows and columns here represent different combinations of the feedback *f*_1_ and *f*_2_ (marked green in Figure 4). A black point superimposed on an ellipse indicates that the trivial state *R*_1_ = *R*_2_ = 0 is also stable.

### 3.4 Discontinuity and multistability at high cooperativity

Crosstalk can not only couple the two responses, as described above, but also restrict the joint responses to just a few distinct states. Figure 6 shows the behavior when the cooperativity *n* is increased to *n* = 2 and *n* = 5. The mesh of Figure 3 breaks up into multiple tight clusters (Figure 6 left column). As a result, there are only three distinct stable states that exist: The on state is characterized by activation of both *R*_1_ and *R*_2_; the half-on state has *R*_2_ activated and *R*_1_ partially activated; the off state has no activation of either *R*_1_ or *R*_2_. Due to the asymmetric, hierarchical positioning of the pathways, there is no fourth state where *R*_1_ is active while *R*_2_ is inactive.

**Figure 6:**
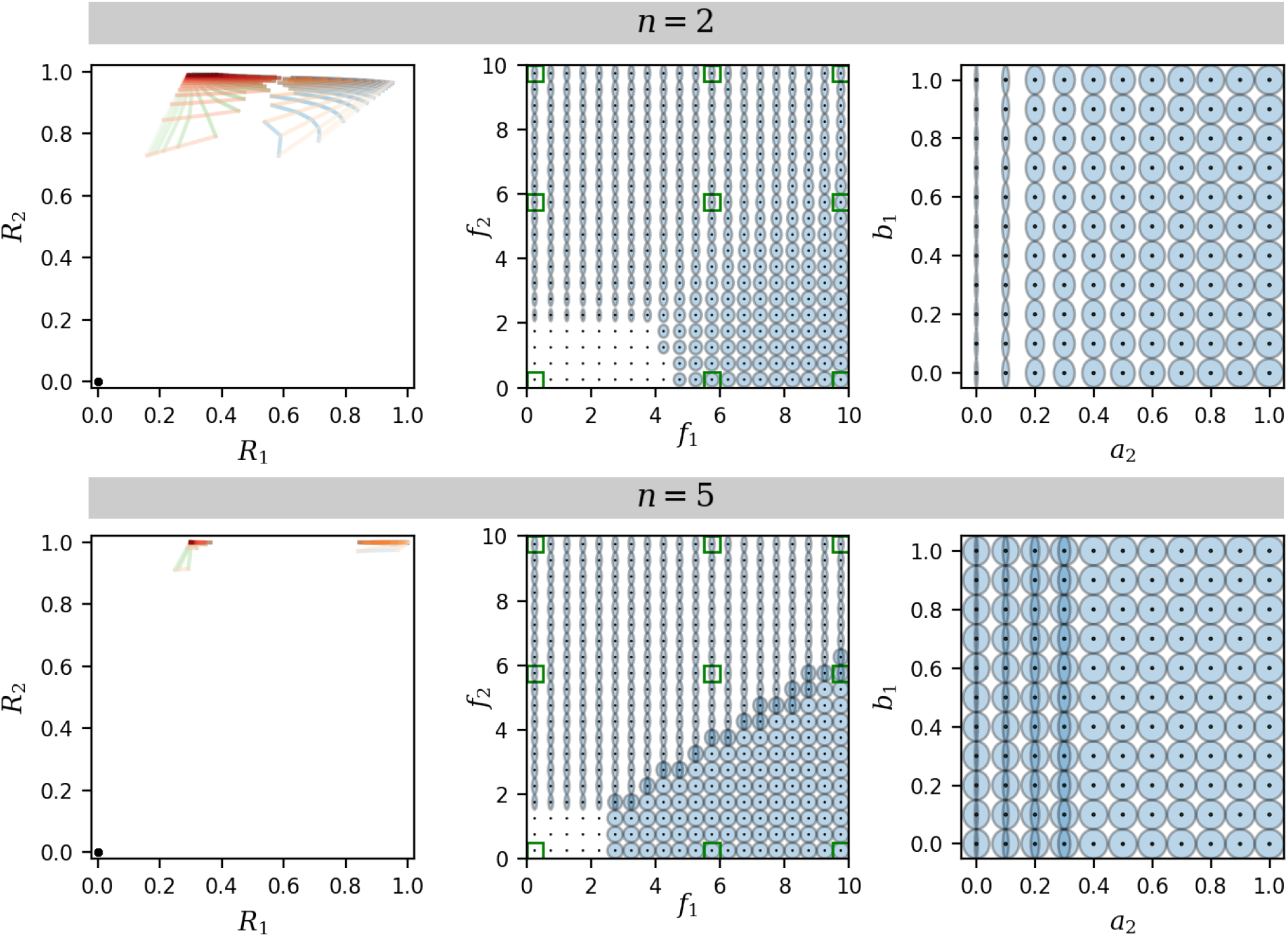
Examples of mesh and ellipse plots for higher cooperativity *n* = 2 and *n* = 5 (See Figures S2 & S3 for detailed plots for *n* = 5). Left column: *a*_2_ = *b*_1_ = 0.3, to be compared with the center panel of Figure 3. Middle column: *a*_2_ = *b*_1_ = 0.3, to be compared with the center panel of Figure 4. Right column: *f*_1_ = 9.8 and *f*_2_ = 5.8, to be compared with the mid-right panel of Figure 5. All panels have *a*_1_ = *b*_2_ = 1.

As can be seen from Figure 6 (middle column), at high cooperativity the responses *R*_1_ and *R*_2_ no longer change smoothly with the feedback strengths *f*_1_ and *f*_2_, but switch discontinuously along certain boundaries in the (*f*_1_, *f*_2_) parameter space (see also Figure S2 for *n* = 5). In the limit *n* → ∞, the phase diagram of Figure 7 is obtained (see Appendix A.3), where each region of the parameter space permits different types of solutions to Eq. (2). In Region I, both feedbacks *f*_1_ and *f*_2_ are too small to activate a response, allowing only the trivial solution with *R*_1_, *R*_2_ both off. In Region II, with high *f*_2_ and relatively low *f*_1_, there exists an additional half-on state, with the upstream *R*_2_ fully activated and the downstream *R*_1_ only partially active. In Region IV, *f*_1_ and *f*_2_ are both sufficiently large to allow simultaneous activation of both responses *R*_1_ and *R*_2_, i.e., a fully on state instead of half-on. Between regions II and IV is region III, which allows both the half-on state and the fully on state, in addition to the off state.

**Figure 7:**
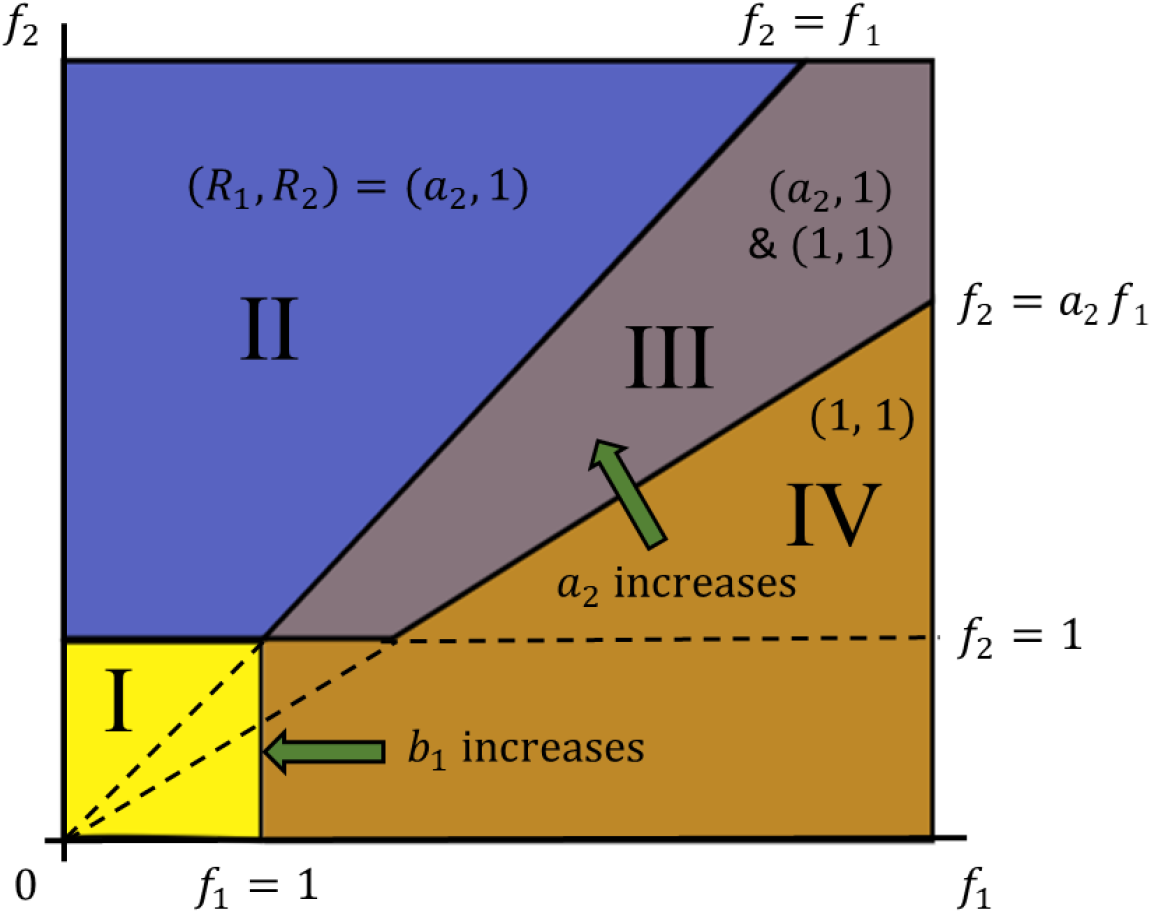
Phase diagram showing which states of (*R*_1_, *R*_2_) are permitted for different combinations of *f*_1_ and *f*_2_ values (in addition to the trivial solution *R*_1_ = *R*_2_ = 0), in the limit of high cooperativity *n* → ∞. (I) there is only the trivial solution - *R*_1_ and *R*_2_ both off; (II) *R*_2_ is on, *R*_1_ is partially on (proportional to *a*_2_); (III) *R*_2_ is on, *R*_1_ can be either fully or partially on; (IV) both *R*_1_ and *R*_2_ are fully on.

Importantly, changes in the crosstalk strengths cause the boundaries in the phase diagram to move, because crosstalk allows the upstream and downstream feedback loops to activate each other. The sloped boundary between Region III and IV depends on the crosstalk parameter *a*_2_: as *a*_2_ increases, this boundary rotates counterclockwise around the origin, expanding Region IV and reducing Region III. In addition, the value of *R*_1_ in Region II & III also increases with *a*_2_. Similarly, the vertical boundary between Regions IV and I depends on *b*_1_: if *b*_1_ decreases to 0, this boundary moves all the way to the right, removing the small-*f*_2_ portion of Region IV.

## 4 Discussion

The model studied here does not contain all the ingredients of any quorum sensing pathway and is not intended as a completely faithful representation of *V. fischeri* LuxRI and AinRS pathways. It does however capture several common properties of quorum sensing networks: (1) multiple signals and cognate receptors drive multiple regulated outputs; (2) these pathways are functionally linked (sequentially in *V. fischeri*); (3) the pathways crosstalk through the interaction of signals with non-cognate receptors; (4) both pathways are regulated with positive feedback. The model offers insight into how the strength of crosstalk interacts with these architectural properties to alter the behavior of a quorum sensing system.

### 4.1 The meaning of crosstalk

Quorum sensing pathways frequently employ multiple ligand-receptor pairs with limited binding specificity, where the “promiscuity” of this binding is a highly tunable or evolvable property [19, 4, 11]. Previous authors have investigated behavior of signaling pathways subject to this and other mechanisms of crosstalk; these include weak selectivity in ligand affinities at a cell surface receptor [16], multiple ligand-receptor channels that merge to control a single regulated output [13, 24], or a receptor that has multiple sensing states and outputs that are associated with binding of different ligands [14].

The ubiquity and tunability of crosstalk in quorum sensing systems raise the question of how even weak crosstalk may enhance the function of a multi-signal system. For example, a limited amount of crosstalk between two signaling pathways can in principle enhance the ability to measure the input signal concentrations [16]. It may also provide some benefit in suppressing early response [25]. Therefore, rather than consider a mechanistic model that embeds strong crosstalk into the topology, we study a model where distinct signaling pathways are coupled through tunable crosstalk parameters: crosstalk strength can range from a perturbation that weakly couples the two pathways to a strong link that drives two regulated outputs in tandem.

### 4.2 Crosstalk through cross-binding or cross-activation

Our analysis highlights the importance of distinguishing between two different aspects of crosstalk that occurs when a receptor interacts with its non-cognate ligand (signal): One is cross-affinity, or the lack of specificity in binding, of a signal by the non-cognate receptor (characterized by parameter *b* in Eq. 1c). The other is the ability of the resulting non-cognate complex to cross-activate the regulated pathway (characterized by parameter *a*). If the receptor bound by the non-cognate signal is ineffective at promoting transcription, the result is competitive inhibition of the receptor. Mathematically, in Eq. (1c), the inhibition is due to *S*_2_ appearing in the denominator of the expression for *R*_1_. Thus, cross-binding allows the excitatory signal of one channel to inhibit the non-cognate response. On the other hand, if the receptor bound by a non-cognate signal can still promote transcription to some level, then there is cross-activation of the response, especially when the cognate signal is absent. Mathematically, when *S*_1_ = 0, Eq. (1c) allows *R*_1_ to be activated by signal *S*_2_, although at a lower saturating level governed by *a*_2_ (see Appendix A.3). Thus, the activation (*a*) and binding (*b*) components of crosstalk allow a response to be either activated or inhibited respectively by the non-cognate signal, when its own cognate signal is weak.

The dual effect of the non-cognate signal is observed in *V. fischeri*, where both C8-HSL and 3OC6-HSL are able to form an activating complex with LuxR. Because the C8-HSL-LuxR complex is less efficient at inducing *lux*, crosstalk from the *ainS/R* pathway inhibits the activating effect of 3OC6-HSL during the growth of a colony [26, 27]. Although the C8-HSL autoinducer can stimulate luminescence at low concentrations of 3OC6-HSL [25], at high 3OC6-HSL concentrations, the addition of C8-HSL reduces luminescence through the competition effect [28].

### 4.3 Weak crosstalk expands dynamic range

The extreme case of *V. harveyi*, where two signals merge to drive a single regulated output, is an example of two dimensions of signal input leading to a one-dimensional regulatory output. In *V. fischeri* the strength of crosstalk is evidently tunable between strains, as the HSL specificity of the LuxR receptor is strain-dependent. Further, both the *lux* regulatory region and the AinS/AinR system exhibit much greater sequence divergence between strain isolates of *V. fischeri* than is typical of the rest of the genome [29]. These findings suggest that the quorum sensing pathways in *V. fischeri* are under strong, strain-dependent selection pressures with consequences for *lux* control and associated crosstalk.

What benefits can different strains gain by tuning weak crosstalk interactions between the two pathways? Figure 3 shows graphically how crosstalk strength modulates the space of system outputs. A sensing system with two fully independent outputs such as *R*_1_ and *R*_2_ has in principle a two-dimensional space of outputs. In the limit of strong crosstalk these two independent outputs are collapsed to fall along the same, one-dimensional arc. Thus, fine tuning of the strength of crosstalk between the signal paths can define the dimensionality, shape, and extent of the response region in Figure 3, tuning the dual output response to *f*_1_ and *f*_2_ along a continuum from orthogonality to tandem or ‘funneled’ control. Funneled control may be beneficial when there is a risk that quorum interference is removing one signal from the environment, so that “OR” sensing is desirable; orthogonality will be advantageous when multiple phenotypic behaviors need to be controlled by the dual-signal system. Crosstalk allows some compromise between these two limiting behaviors.

### 4.4 Role of feedback

With both nonlinearity and feedback present in the model, multistability of output can be expected, especially in the limit of high cooperativity. Accordingly we observe multistable states in our model in several regions of the parameter space. Although multistability normally requires *n* ≥ 2, in the presence of crosstalk there is multistability even for *n* = 1, such as when *b*_1_ *>* 0 and *f*_1_ is high but *f*_2_ is low (Figure 5 bottom right). This is because the coupling between the two feedback loops results in stronger nonlinearity than in a single feedback loop, allowing multistability (see Appendix A.3). Experiments however find few clear examples of multistability in quorum sensing. With some exceptions (usually based on synthetic or ‘rewired’ pathways [30, 31, 32]), multistability is rarely observed in Gram negative quorum systems. Generally the highly diffusible HSL signals, together with positive feedback synthesis, act intercellularly to lock the entire population into an on-state. Because of extracellular accumulation of autoinducer, the off-state becomes increasingly unfavorable or unlikely compared to the on-state. Multistability in quorum sensing circuits is more evident when the signal feedback at the individual cell level is strengthened, either by circuit redesign [31] or because the signal is able to act intracellularly, allowing isolated cells to autoactivate [33].

Another factor that makes it difficult to observe multistability in individual cells is that experiments often fix the extracellular signal concentrations at defined levels, or use strains in which feedback has been broken by deletion of the signal synthases. Thus, the difference between the multistability in experimental conditions and in our model highlights the important distinction between keeping the signals constant and letting them “float” according to the feedback. To observe multistability one must allow the signal levels to float either upward or downward on the route to steady state, an atypical experimental condition. Instead, under typical conditions, multistability is suppressed and noise in gene expression is a much more significant source of heterogeneity [34, 35].

### 4.5 New motifs: “jump-start” and “push-start”

Our results show that crosstalk can allow either of the feedback loops – upstream or downstream – to activate the other loop, via separate mechanisms driven by *a*_2_ and *b*_1_ respectively. As illustrated schematically in Figure 8A, the first mechanism is engaged when the feedback strength *f*_1_ is low but *f*_2_ is high (Region II in Figure 7). A large *f*_2_ turns on the *S*_2_-*R*_2_ feedback loop, but without crosstalk the *R*_1_ response is off due to a small *f*_1_. Strengthening the downstream-directed crosstalk *a*_2_ allows the upstream *S*_2_-*R*_2_ feedback loop to also activate the downstream *R*_1_ response (Figure 8A). This is roughly analogous to the “jump-start” of a combustion engine, where an upstream system consisting of a battery, alternator and starter motor is mechanically coupled to a downstream system consisting of the combustion engine and flywheel: activating the upstream branch by energizing the starter system turns the motor which then starts the combustion engine.

**Figure 8:**
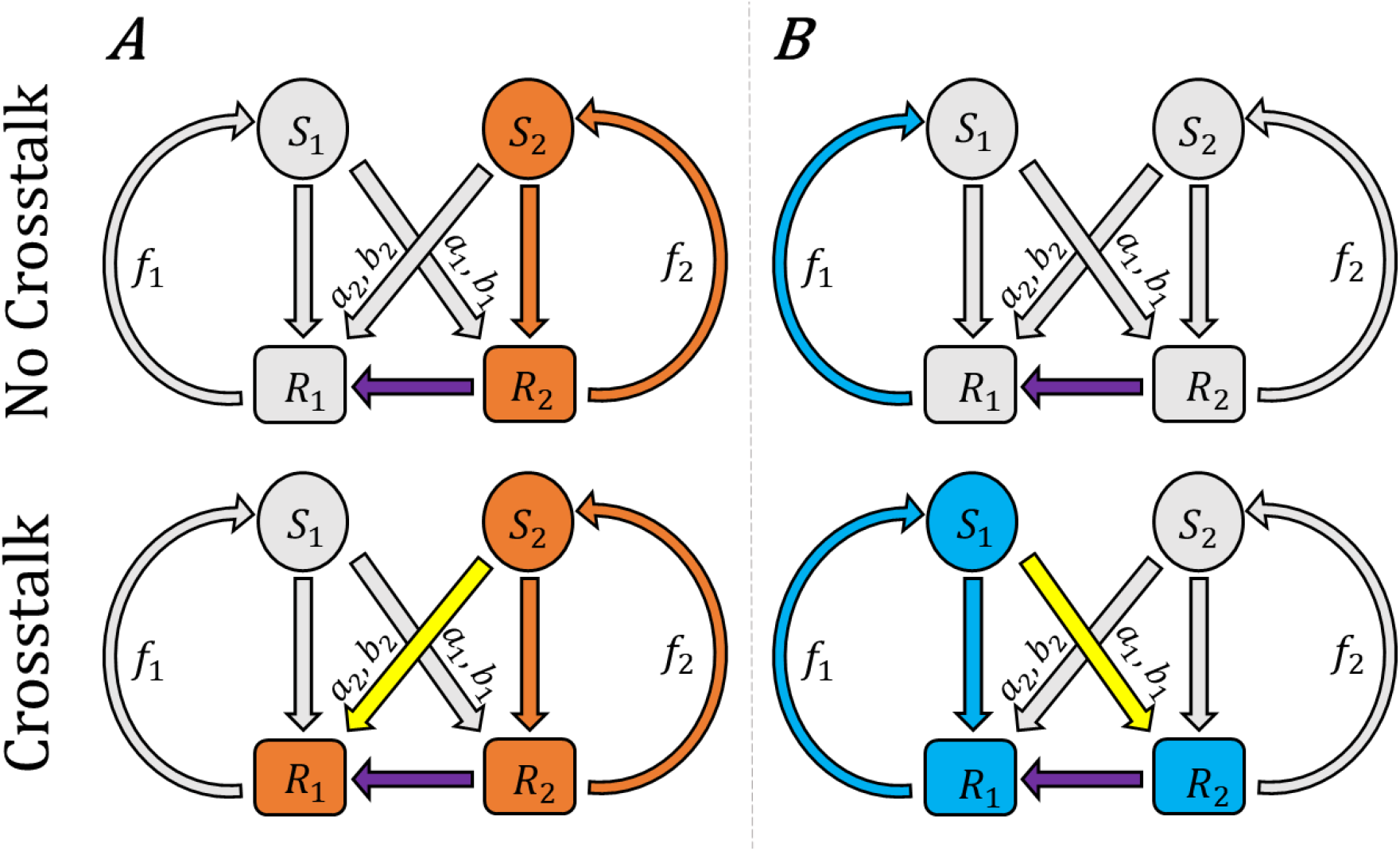
Crosstalk provides two new mechanisms for activating both pathways without requiring strong feedback in both. In the “jump start” scenario (left), even when the feedback *f*_1_ is weak but *f*_2_ is strong, the downstream-directed crosstalk (*a*_2_, *b*_2_) allows the *S*_2_-*R*_2_ pathway to activate and drive the downstream *R*_1_ response. In the “push start” scenario (right), when the feedback *f*_2_ is weak but *f*_1_ is strong, the upstream-directed crosstalk (*a*_1_, *b*_1_) allows the *R*_2_ response to be driven by the *S*_1_-*R*_1_ pathway, activating both *R*_1_ and *R*_2_.

The second mechanism, shown in Figure 8B, applies when *f*_1_ is large but *f*_2_ is small (lower part of Region IV in Figure 7). In the absence of crosstalk, even a large *f*_1_ cannot turn on *R*_1_, because it depends on the upstream response *R*_2_ that is off due to a small *f*_2_. However, sufficient upstream-directed crosstalk *b*_1_ can allow the downstream *R*_1_-*S*_1_ feedback to drive the upstream *R*_2_ and activate both *R*_1_ and *R*_2_ (Figure 8B). This has a rough analogy in the “push-start” of a combustion engine with a dead starter battery: the mechanical coupling from the engine crankshaft to the upstream battery/alternator system allows an energy input at the wheels to turn the engine, which then turns the alternator, replacing the role of the battery and activating the downstream (engine) and upstream (alternator/battery) systems.

These two mechanisms can be thought of as new regulatory motifs that could be embedded within larger gene regulatory networks. The essence is that two feedback loops, linked by crosstalk, are positioned upstream and downstream from each other. Crosstalk between the feedback loops allows activation through the jump-start (upstream feedback loop activating downstream response) and push-start (downstream feedback loop activating both responses) behaviors. In addition to these motifs (where we set *a*_1_ = *b*_2_ = 1), there are other interesting behaviors when all the crosstalk parameters are considered, as summarized in Table 1. In particular, we have the opposite of the jump-start and push-start, which could be called “jump-stop” and “push-stop”, where crosstalk from one branch can inhibit the function of the other branch.

**Table 1:** Modes of activation and deactivation through crosstalk between two coupled quorum sensing pathways. The “jump start” and “push start” modes activate the system as illustrated in Figure 8, whereas the “jump stop” and “push stop” modes shut down the sysytem. The *f*_1_, *f*_2_ columns show the feedback conditions required for each mode, while the *a*_1_-*b*_2_ columns show how the crosstalk parameters must be configured, to generate the outputs shown in the *R*_1_, *R*_2_ columns. A dash means the parameter does not strongly impact the behavior.

| Type | $f_1$ | $f_2$ | $a_1$ | $a_2$ | $b_1$ | $b_2$ | $R_1$ | $R_2$ |
| --- | --- | --- | --- | --- | --- | --- | --- | --- |
| Jump Start | Low | High | 1 | ↑ | — | 1 | ↑ | 1 |
| Push Start | High | Low | 1 | — | ↑ | 1 | ↑ | ↑ |
| Jump Stop | High | High | — | 0 | — | ↑ | ↓ | 1 |
| Push Stop | High | High | 0 | — | ↑ | — | ↓ | ↓ |

## 5 Conclusion

Although crosstalk in engineering contexts refers to an undesired leakage of information between separate communication channels, in the context of biological sensing it can provide additional mechanisms for the control or activation of coupled feedback systems that are ubiquitous in quorum sensing pathways. In our analysis crosstalk appears to provide a route for switching individual feedback circuits on or off without relying entirely on extracellular signal concentrations as in typical interpretations of quorum sensing. Our findings are based on analyzing the equilibrium states of the feedback circuits that are coupled through crosstalk. How crosstalk affects the kinetics of the system as it approaches the equilibrium is likely an important component of its biological role, which remains to be studied.

The variability of crosstalk strength across different quorum sensing systems and even across strains of the same bacterial species suggests that the new mechanisms for activating the feedback circuits can be exploited through evolutionary tuning of the crosstalk strength. For example, crosstalk could provide a form of redundancy so that a pathway can still be activated when the signal is being inhibited, such as by quorum interference: If a sabotaging species removes the signal *S*_1_ from the environment, or creates an interfering signal *S*_3_ that saturates the *S*_1_ receptor, the jump-start mechanism may ensure that the downstream response *R*_1_ can still be activated. Likewise, the push-start mechanism may protect against external interference with the signal *S*_2_ of the upstream response *R*_2_.

It may also be possible that crosstalk strengths could be tuned on short timescales by cellular processes. For example, the cross-binding strength *b* could be affected by allosteric interactions with modifier proteins, while the cross-activation efficiency *a* could be controlled by other ligands or post-translational regulation. If crosstalk was variable in real time, instead of over evolutionary timescales, then it could be a very significant mechanism for control. It would be interesting if experiments could show that crosstalk strengths are variable within the same species and under different environmental conditions, allowing jump-start and push-start in real time. This could even lead to community-level phenomena, such as one species triggering the quorum sensing pathway of another by tuning their crosstalk strengths, i.e., an “interspecies jump-start”.

## Acknowledgements

Funding support from National Science Foundation award MCB 1715981 is acknowledged.

## A Methods

### A.1 Formulating the Mathematical Model

We consider a quorum sensing system in which two autoinducer signals drive the activity of two largely distinct, but coupled, regulatory pathways. In a simplified picture, where the two pathways operate without crosstalk, the autoinducer binds to a cognate receptor and the bound complex promotes the expression of a corresponding set of genes, including one that encodes the autoinducer itself. The specific mechanisms of signal transduction are variable, and may involve an intracellular receptor that binds the signal to form a transcriptional activator, or a membrane bound receptor that controls a phosphorylation cascade.

We think of the signals *S*_*i*_ as the concentrations of the two autoinducer species, and the responses *R*_*i*_ as the expression levels of the corresponding quorum regulated genes. We model the gene expression level as a function of the autoinducer concentration using a binding-equilibrium expression that resembles the Michaelis-Menten equation,

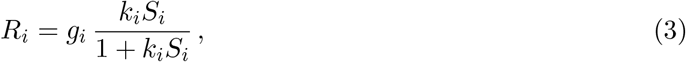

where *g*_*i*_ is the maximum level at saturation and *k*_*i*_ represents the binding affinity of the autoinducer. For each pathway the expression level *R*_*i*_ of quorum sensing genes determines the production of the corresponding autoinducer *S*_*i*_, through positive regulatory feedback. For simplicity, we assume that:

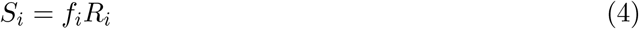

where *f*_*i*_ is the strength of feedback and depends on the diffusion and degradation of the autoinducer molecules.

Crosstalk between the two quorum sensing pathways occurs when the autoinducer of one pathway can modulate gene expression in the other pathway. For example, a non-cognate autoinducer may bind to the receptor with some affinity, producing a resultant complex that promotes gene expression with some efficiency. Thus, the gene expression level will depend on the concentration of both autoinducers. We model such dependence by modifying Eq. (3) to:

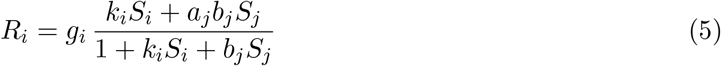

Here *j* is the label for the non-cognate signal – the *b*_*j*_*S*_*j*_ term in the denominator represents competitive binding by the non-cognate autoinducer, and the *a*_*j*_*b*_*j*_*S*_*j*_ term in the numerator represents cross-activation by the non-cognate complex. The parameter *b*_*j*_ represents the binding affinity of the non-cognate autoinducer to the receptor, and *a*_*j*_ represents the promotion efficiency of the non-cognate complex. This form of dependence on multiple signals is fairly general as it can be derived for different, common quorum sensing system architectures [24, 25]. Detailed derivation of these equations for the specific pathways in *V. fischeri* is described in the Supplementary Material.

We now incorporate some more details of the circuit that is present in the *Vibrio fischeri* example. In that system, pathway 2 is upstream of pathway 1 (Figure 1A), so that the expression level of *R*_1_ depends on that of *R*_2_ (Figure 1B). This is modeled by simply making *R*_1_ proportional to *R*_2_,

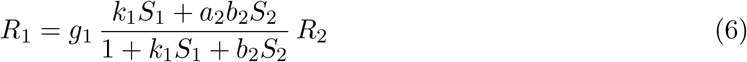

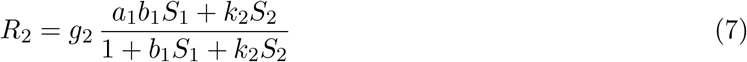

We may remove the parameters *g*_*i*_ and *k*_*i*_ by rescaling *S*_1_ → *S*_1_*/k*_1_, *S*_2_ → *S*_2_*/k*_2_, *R*_1_ → *g*_1_*g*_2_*R*_1_, and *R*_2_ → *g*_2_*R*_2_, and redefining parameters *b*_1_ → *b*_1_*k*_1_, *b*_2_ → *b*_2_*k*_2_. After such rescaling the equations finally become:

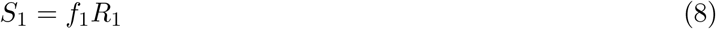

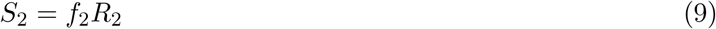

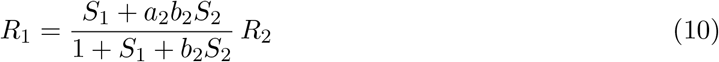

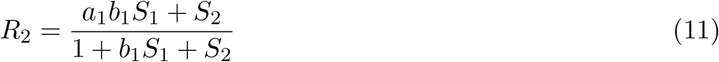

### A.2 Numerical methods for finding solutions

To find the solutions to Eqs. (8–11), we consider a system of differential equations whose equilibrium states are the solutions to those equations above. The differential equations we use are:

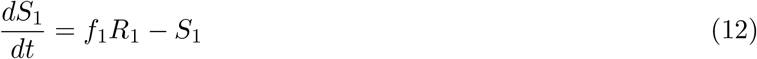

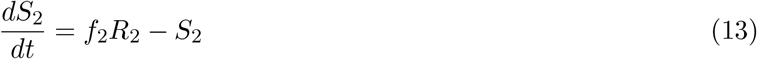

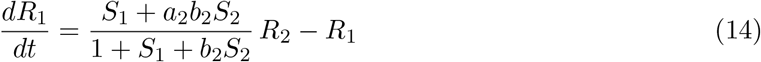

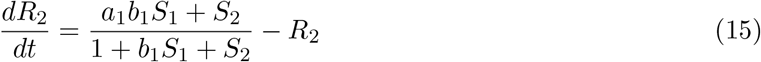

For each set of parameter values, we integrate these equations using the SciPy function solve IVP() for 500 time units, starting from random initial values. This should bring the variables sufficiently close to a local equilibrium. We then refine the result using SciPy’s root-finding function scipy.optimize.root(), with the previous result as the initial guess. Alongside this, we calculate the Jacobian matrix of our system of equations to verify that the equilibrium that we have found is stable. In order to find all potential solutions for a given set of parameters, we repeat this process 100 times with different random initial values and eliminate any duplicate solutions. To generate the figures in the main text, we scan over the parameter space and apply the above procedure at every grid point in the parameter space.

### A.3 Analytic results at *n* → ∞

In the limit *n* → ∞, the solutions to Eqs. (2) can be found using the following arguments:

a. If both *R*_1_ and *R*_2_ are small, such that *f*_1_*R*_1_ < 1 and *f*_2_*R*_2_ < 1, then the right-hand side (RHS) would lead to *R*_1_, *R*_2_ → 0. Indeed, *R*_1_ = *R*_2_ = 0 is always a solution.
b. If *f*_2_*R*_2_ *>* 1 and *f*_2_*R*_2_ *> f*_1_*R*_1_, then the RHS gives *R*_2_ = 1 and *R*_1_ = *a*_2_. To be consistent, we need *f*_2_ *>* 1 and *f*_2_ *> a*_2_*f*_1_, which corresponds to Regions II and III in Figure 7.
c. If *f*_1_*R*_1_ *> f*_2_*R*_2_ *>* 1, then the RHS gives *R*_1_ = *R*_2_ = 1. To be consistent, we need *f*_1_ *> f*_2_ *>* 1, which corresponds to Region III and part of Region IV in Figure 7.
d. If *f*_1_*R*_1_ *>* 1 *> f*_2_*R*_2_, then the result depends on *b*_1_. For *b*_1_ *>* 0, the RHS gives *R*_1_ = *R*_2_ = 1 as in (c), which corresponds to the *f*_1_ *>* 1 *> f*_2_ part of Region IV in Figure 7. But for *b*_1_ = 0, the RHS gives *R*_1_ = *R*_2_ = 0, which means this part is merged into Region I.

Taken together, the above arguments imply that:

i. In Region I, the only solution is (*R*_1_, *R*_2_) = (0, 0).
ii. In Region II, both solutions (0, 0) and (*a*_2_, 1) are possible.
iii. In Region III, three solutions are possible, (0, 0), (*a*_2_, 1), and (1, 1).
iv. In Region IV, two solutions are possible, (0, 0) and (1, 1).

These different regions are shown in Figure 7.

In some regions of interest, approximate solutions can be obtained to understand the behavior of the system. In the case where *f*_1_ is small (left part of Region II), we can approximate that *S*_1_ ≈ 0. This allows us to ignore the effect of *S*_1_ and simplify the equations to:

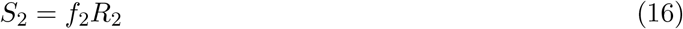

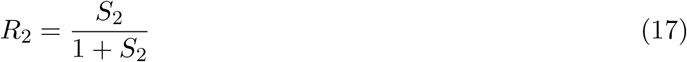

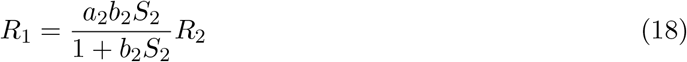

In this region, the *R*_1_ response is completely controlled by the crosstalk coming from the *S*_2_-*R*_2_ feedback loop, which is the “jump-start” scenario. The activation level of *R*_1_ is controlled by the parameter *a*_2_, while its dependence on *b*_2_ is weak as long as the feedback *f*_2_ is strong enough to highly express *S*_2_ (i.e., *f*_2_ > 1*/b*_2_).

Similarly, when *f*_2_ is small (lower part of Region IV), we can ignore *S*_2_ and simplify the equations to:

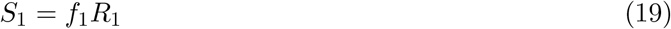

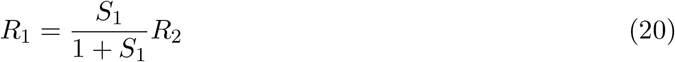

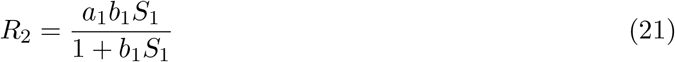

In this case, when *a*_1_ and *b*_1_ are not too small, there is a feedback loop *S*_1_-*R*_2_-*R*_1_ that can activate both responses *R*_1_ and *R*_2_ to a level controlled by *a*_1_, which is the “push-start” scenario. For *a*_1_ = 1, both responses can be fully activated as long as the feedback *f*_1_ is strong enough (*f*_1_ *>* 1*/b*_1_). In other words, the responses can be turned on by tuning the parameter *b*_1_ above a threshold ∼ 1*/f*_1_. Note that if we eliminate *R*_2_ from Eqs. (20,21), then *R*_1_ as a function of *S*_1_ is more nonlinear than a Hill’s function with *n* = 1.

## S1 Supplementary Text

### S1.1 Derivation of crosstalk equations based on the specific *V. fischeri* circuit

The signaling system with two coupled pathways modeled in the main text is based loosely on the *luxIR* and *ainRS* quorum sensing pathways in *V. fischeri*. Here we go into some biochemical detail to motivate equation (5), which describes the dependence of the response variables *R*_*i*_ on both signals. In the case of *V. fischeri*, the two signals *S*_1_ and *S*_2_ may represent the autoinducers 3OC6-HSL and C8-HSL respectively, and the two responses *R*_1_ and *R*_2_ may be analogous to the expression of the *luxI* and *ainS* genes that encode the synthases of the autoinducers respectively. For pathway 1, the 3OC6-HSL binds reversibly to the transcription factor *LuxR* with an association constant *k*_1_. In addition, *LuxR* has a cross-interaction with C8-HSL, with an association constant *b*_2_. Let [*LuxR*_0_] and [*LuxR*] be the concentration of total and unbound *LuxR*, then:

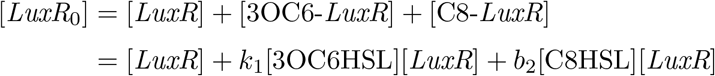

Both complexes 3OC6-*LuxR* and C8-*LuxR* can activate the transcription of the *luxI* gene, with different efficiencies *a*_1_ and *a*_2_. If the promoter is not saturated, the transcription rate will be linear in both complexes:

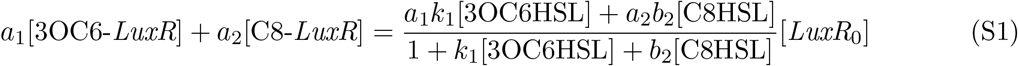

The total amount of *LuxR* depends on the expression of *LitR*, which is a transcription factor for the *luxR* gene, as well as for the *ainS* gene in the other (upstream) quorum sensing pathway. Therefore, we assume that [*LuxR*_0_] ∝ *R*_2_. Writing [3OC6] as *S*_1_ and [C8] as *S*_2_, we have:

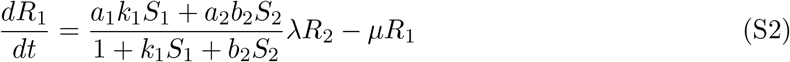

where *µ* is the loss or degradation rate. At equilibrium, we find:

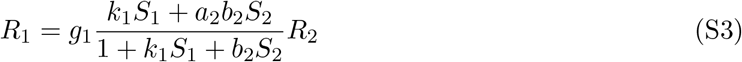

where we defined *g*_1_ ≡ *a*_1_*λ/µ* and redefined *a*_2_ → *a*_2_*/a*_1_.

If we assume a certain degree of multi-merization in the binding of HSL (either 3OC6 or C8) to *LuxR*, i.e.,

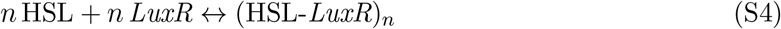

then the above equation will become:

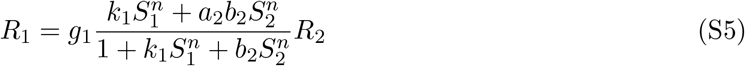

Experimental estimates of the coefficient *n* range roughly between 1– 2 [25].

For pathway 2, the C8-HSL binds to a transmembrane receptor *AinR*, which is a histidine kinase that triggers a signal transduction through a phosphorelay protein *LuxU*. Binding of the HSL to the receptor eventually leads to the increase of the *LitR* level, which promotes the transcription of the *ainS* gene. It has been shown that the 3OC6-HSL can also promote the expression of *LitR*, although the mechanism has not been fully determined. In a homologous quorum sensing pathway in *V. harveyi, LuxU* can be phosphorylated by multiple receptors; the dependence of the shared output variable on the input from each autoinducer was modeled using a similar form to our Eq. (5) [24] (see Eq. (4) therein).

## S2 Supplementary Figures

**Figure S1:**
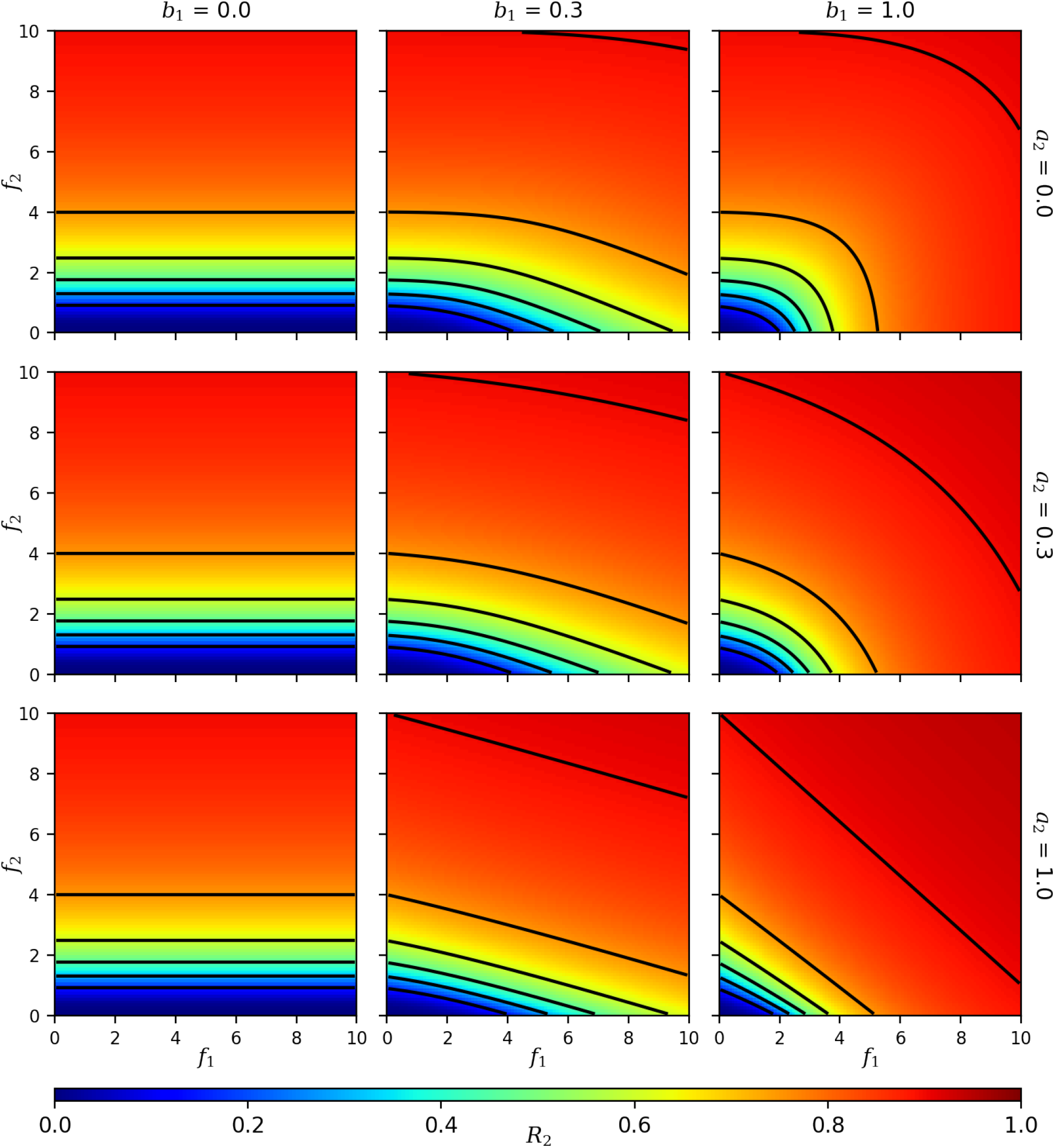
Heat maps showing the value of *R*_2_ as a function of the feedback strengths *f*_1_ and *f*_2_ (compare with Figure 2 for *R*_1_). Rows correspond to different values of the downstream-directed crosstalk activation strength *a*_2_, whereas columns correspond to values of the upstream-directed crosstalk binding strength *b*_1_ (all panels have *a*_1_ = *b*_2_ = 1, *n* = 1). Black lines are contours of constant *R*_2_.

**Figure S2:**
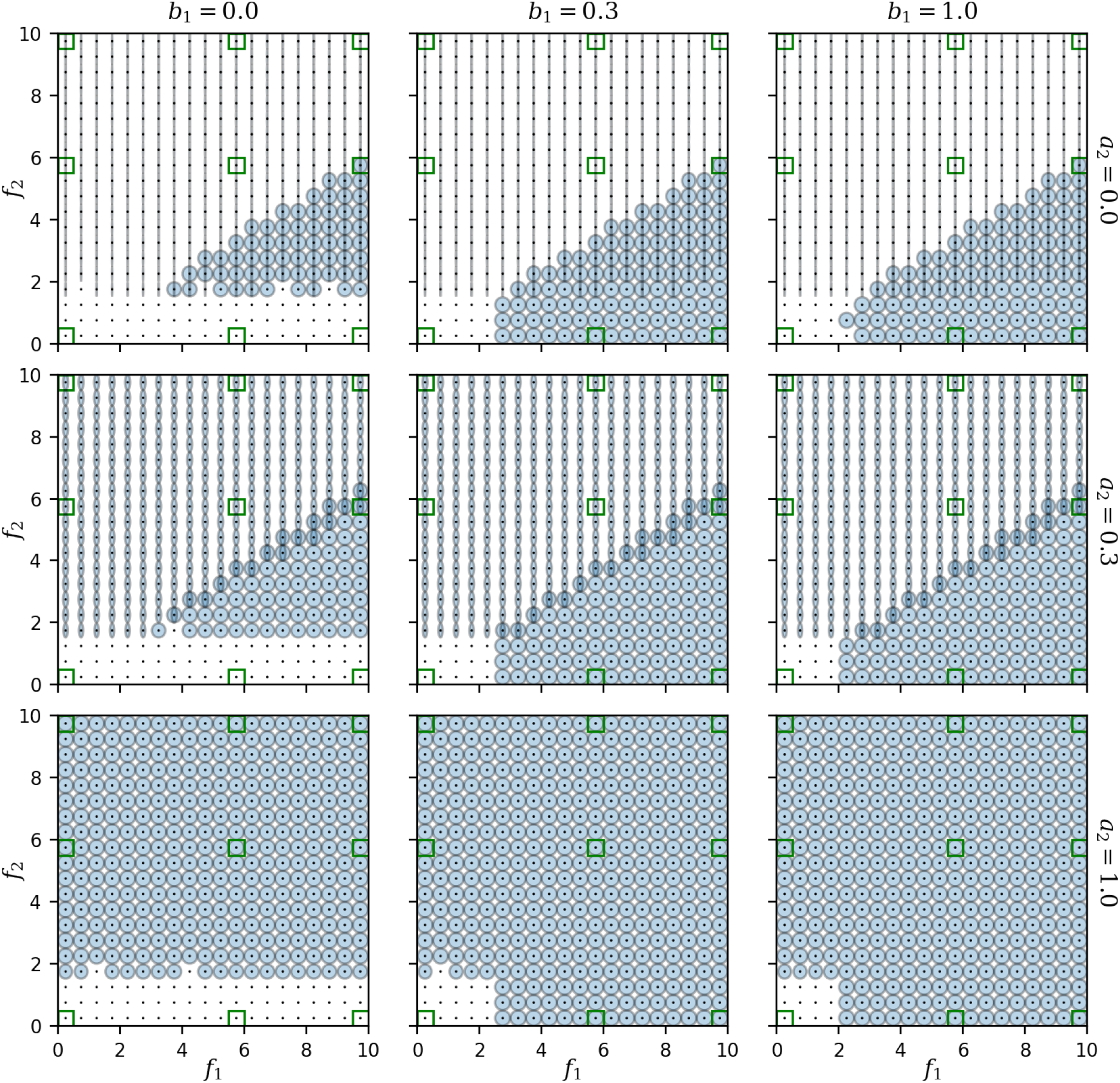
“Ellipse plots” showing the responses *R*_1_ and *R*_2_ as functions of the feedback strengths *f*_1_ and *f*_2_ for high cooperativity *n* = 5 (compare with Figure 4 for *n* = 1). The width and height of each ellipse represent the values of *R*_1_ and *R*_2_, respectively, at a given point in the (*f*_1_, *f*_2_) plane. Multiple ellipses overlaid on top of one another represent multiple stable solutions; a black point represents a vanishing ellipse with *R*_1_ = *R*_2_ = 0. Rows and columns represent different values of the crosstalk parameters *a*_2_ and *b*_1_, respectively (with *a*_1_ = *b*_2_ = 1). The *f*_1_ and *f*_2_ values marked in green are shown in Figure S3.

**Figure S3:**
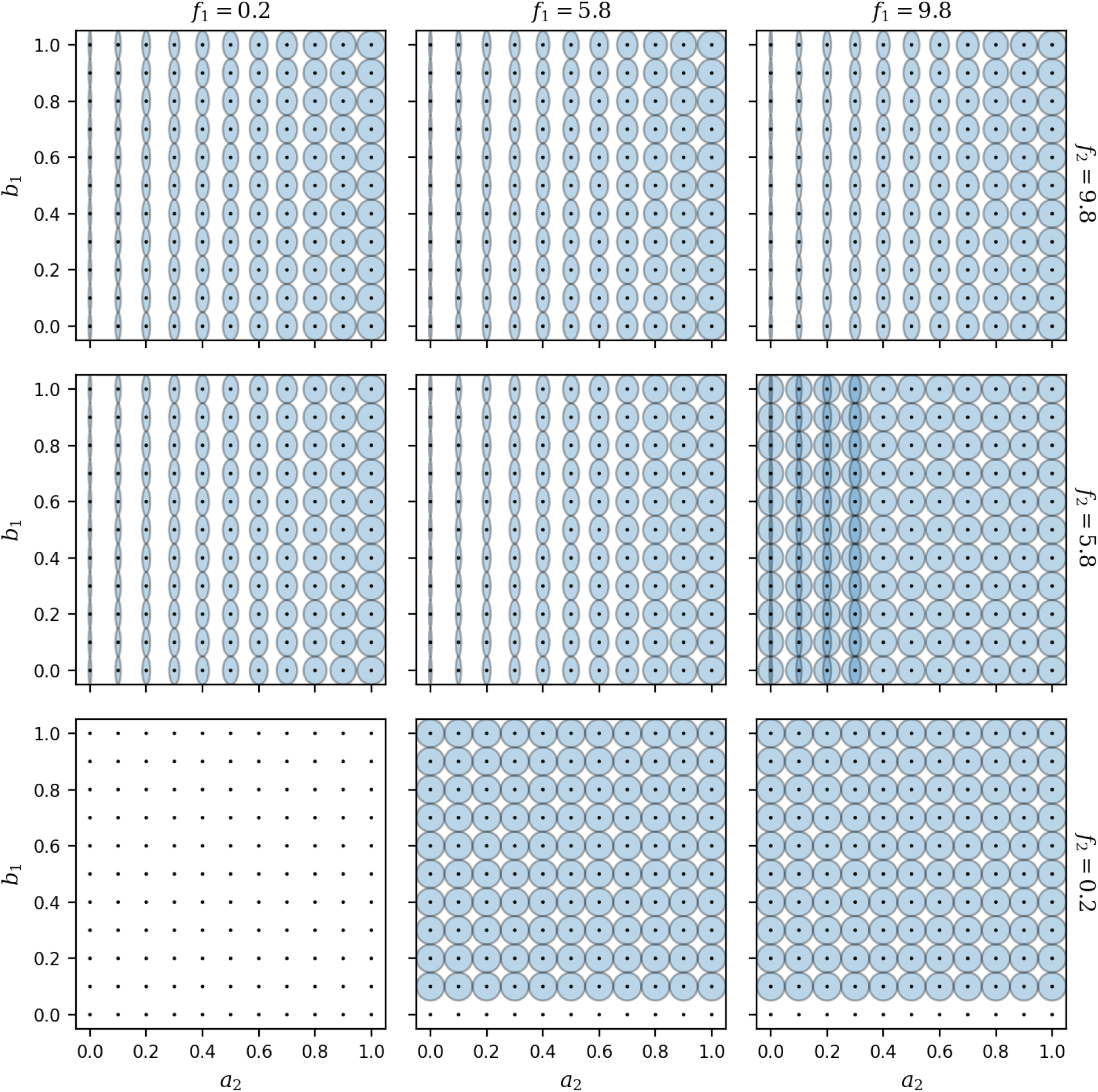
Dependence of the responses *R*_1_ and *R*_2_ on the crosstalk parameters *a*_2_ and *b*_1_ for high cooperativity *n* = 5 (compare with Figure 5 for *n* = 1). With the same parameters as in Figure S2, the width and height of each ellipse represent the values of *R*_1_ and *R*_2_, respectively. Rows and columns here represent different combinations of the feedback *f*_1_ and *f*_2_ (marked green in Figure S2). Multiple ellipses overlaid on top of one another represent multiple stable solutions; a black point represents a vanishing ellipse with *R*_1_ = *R*_2_ = 0.

## References

[1] K. Papenfort and B.L. Bassler. Quorum sensing signal-response systems in gram-negative bacteria. Nature Reviews Microbiology, 14:576–588, 2016.

[2] S. Mukherjee and B.L. Bassler. Bacterial quorum sensing in complex and dynamically changing environments. Nature Reviews Microbiology, 17:371–382, 2019.

[3] Nitzan Aframian and Avigdor Eldar. A bacterial tower of babel: Quorum-sensing signaling diversity and its evolution. Annual Review of Microbiology, 74(1):587–606, 2020.

[4] Lisa A. Hawver, Sarah A. Jung, and Wai Leung Ng. Specificity and complexity in bacterial quorum-sensing systems. FEMS Microbiology Reviews, 40(5):738–752, 2016.

[5] Christopher M. Waters and Bonnie L. Bassler. Quorum sensing: Cell-to-cell communication in bacteria. Annual Review of Cell and Developmental Biology, 21(1):319–346, 2005.

[6] Jessica R. Gooding, Amanda L. May, Kathryn R. Hilliard, and Shawn R. Campagna. Establishing a quantitative definition of quorum sensing provides insight into the information content of the autoinducer signals in Vibrio harveyi and Escherichia coli. Biochemistry, 49(27):5621–5623, 2010.

[7] Claudia Anetzberger, Matthias Reiger, Agnes Fekete, Ursula Schell, Nina Stambrau, Laure Plener, Joachim Kopka, Phillippe Schmitt-Kopplin, Hubert Hilbi, and Kirsten Jung. Autoinducers act as biological timers in Vibrio harveyi. PLOS ONE, 7(10):1–12, 2012.

[8] Daniel M. Cornforth, Roman Popat, Luke McNally, James Gurney, Thomas C. Scott-Phillips, Alasdair Ivens, Stephen P. Diggle, and Sam P. Brown. Combinatorial quorum sensing allows bacteria to resolve their social and physical environment. Proceedings of the National Academy of Sciences, 111(11):4280–4284, 2014.

[9] E. Even-Tov, S. Omer Bendori, X. Valastyan, J.and Ke, S. Pollak, and et al. Social evolution selects for redundancy in bacterial quorum sensing. PLOS Biology, 14:e1002386, 2016.

[10] Pulin Li, Michael B. Elowitz, Allon Klein, and Barbara Treutlein. Communication codes in developmental signaling pathways. Development, 146(12), 2019.

[11] Samantha Wellington and E. Greenberg, E. Peter. Quorum sensing signal selectivity and the potential for interspecies cross talk. mBio, 10(2):1–14, 2019.

[12] Fuqing Wu, David J. Menn, and Xiao Wang. Quorum-sensing crosstalk-driven synthetic circuits: From unimodality to trimodality. Chemistry & Biology, 21(12):1629–1638, 2014.

[13] J.M. Henke and B.L. Bassler. Three parallel quorum-sensing systems regulate gene expression in Vibrio harveyi. Journal of Bacteriology, 186:6902–6914, 2004.

[14] Duncan Kirby, Jeremy Rothschild, Matthew Smart, and Anton Zilman. Pleiotropy enables specific and accurate signaling in the presence of ligand cross talk. Physical Review E, 103:042401, 2021.

[15] Michael T. Laub and Mark Goulian. Specificity in two-component signal transduction pathways. Annual Review of Genetics, 41(1):121–145, 2007.

[16] Martin Carballo-Pacheco, Jonathan Desponds, Tatyana Gavrilchenko, Andreas Mayer, Roshan Prizak, Gautam Reddy, Ilya Nemenman, and Thierry Mora. Receptor crosstalk improves concentration sensing of multiple ligands. Physical Review E, 99(2):22423, 2019.

[17] G. Ostovar, K.L. Naughton, and J.Q. Boedicker. Computation in bacterial communities. Physical Biology, 17:061002, 2020.

[18] Deanna M. Colton, Eric V. Stabb, and Stephen J. Hagen. Modeling analysis of signal sensitivity and specificity by Vibrio fischeri LuxR variants. PLOS ONE, 10(5):1–21, 2015.

[19] PK Grant, N Dalchau, JR Brown, F Federici, TJ Rudge, and B Yordanov. Orthogonal intercellular signaling for programmed spatial behavior. Molecular Systems Biology, 12:1–13, 2016.

[20] Subhash Verma and Tim Miyashiro. Quorum sensing in the squid-vibrio symbiosis. International Journal of Molecular Sciences, 14(8):16386–16401, 2013.

[21] J.H. Kimbrough and E.V. Stabb. Substrate specificity and function of the pheromone receptor ainr in Vibrio fischeri es114. Journal of Bacteriology, 195:5223–5232, 2013.

[22] Tim Miyashiro and Edward G. Ruby. Shedding light on bioluminescence regulation in Vibrio fischeri. Molecular Microbiology, 84(5):795–806, 2012.

[23] A.L. Schaefer, B.L. Hanzelka, A. Eberhard, and E.P. Greenberg. Quorum sensing in Vibrio fischeri : probing autoinducer-luxr interactions with autoinducer analogs. J Bacteriol., 178:2897–901, 1996.

[24] Pankaj Mehta, Sidhartha Goyal, Tao Long, Bonnie L Bassler, and Ned S Wingreen. Information processing and signal integration in bacterial quorum sensing. Molecular Systems Biology, 5(1):325, 2009.

[25] Pablo D Perez, Joel T Weiss, and Stephen J Hagen. Noise and crosstalk in two quorum-sensing inputs of Vibrio fischeri. BMC Systems Biology, 5(1):153, 2011.

[26] Christina Kuttler and Burkhard A. Hense. Interplay of two quorum sensing regulation systems of Vibrio fischeri. Journal of Theoretical Biology, 251(1):167–180, 2008.

[27] Alan Kuo, Sean M. Callahan, and Paul V. Dunlap. Modulation of luminescence operon expression by n-octanoyl-l-homoserine lactone in ainS mutants of Vibrio fischeri. Journal of Bacteriology, 178(4):971 –976, 1996.

[28] Claudia Lupp, Mark Urbanowski, E Peter Greenberg, and Edward G Ruby. The Vibrio fischeri quorum-sensing systems ain and lux sequentially induce luminescence gene expression and are important for persistence in the squid host. Molecular Microbiology, 50(1):319–331, 2003.

[29] Jeffrey L. Bose, Michael S. Wollenberg, Deanna M. Colton, Mark J. Mandel, Alecia N. Septer, Anne K. Dunn, and Eric V. Stabb. Contribution of rapid evolution of the luxR-luxI intergenic region to the diverse bioluminescence outputs of Vibrio fischeri strains isolated from different environments. Applied and Environmental Microbiology, 77(7):2445–2457, 2011.

[30] C. Anetzberger, T. Pirch, and K Jung. Heterogeneity in quorum sensing-regulated bioluminescence of Vibrio harveyi. Molecular Microbiology, 73:267–277, 2009.

[31] Eric L. Haseltine and Frances H. Arnold. Implications of rewiring bacterial quorum sensing. Applied and Environmental Microbiology, 74(2):437–445, 2008.

[32] Joshua W Williams, Xiaohui Cui, Andre Levchenko, and Ann M Stevens. Robust and sensitive control of a quorum-sensing circuit by two interlocked feedback loops. Molecular Systems Biology, 4(1):234, 2008.

[33] S.A.M. Underhill, R.C. Shields, J.R. Kaspar, M. Haider, R.A. Burne, and S.J. Hagen. Intra-cellular signaling by the comRS system in Streptococcus mutans genetic competence. mSphere, 3:e00444–18, 2018.

[34] Bianca Striednig and Hubert Hilbi. Bacterial quorum sensing and phenotypic heterogeneity: how the collective shapes the individual. Trends in Microbiology, 30(4):379–389, 2022.

[35] Vera Bettenworth, Benedikt Steinfeld, Hilke Duin, Katrin Petersen, Wolfgang R. Streit, Ilka Bischofs, and Anke Becker. Phenotypic heterogeneity in bacterial quorum sensing systems. Journal of Molecular Biology, 431(23):4530–4546, 2019.

